# Aging Differentially Remodels the Retina and RPE/Choroid Proteome and Metabolism

**DOI:** 10.64898/2026.10.07.757409

**Authors:** Siyan Zhu, Artjola Puja, Rachel McNeel, Diana Alabdallat, Tuan Ngo, Jianhai Du

**Affiliations:** Department of Ophthalmology and Visual Sciences, West Virginia University, Morgantown, WV 26506; Department of Biochemistry and Molecular Medicine, West Virginia University, Morgantown, WV 26506; Department of Pharmaceutical and Pharmacological Science, West Virginia University, Morgantown, WV 26506; Department of Ophthalmology and Visual Sciences, University of Wisconsin-Madison, Madison, WI 53705, USA; McPherson Eye Research Institute, University of Wisconsin-Madison, Madison, WI 53705, USA

**Keywords:** Aging, metabolism, retina, RPE, quantitative proteomics

## Abstract

**Purpose:** The neural retina and retinal pigment epithelium/choroid (RPE/Cho) are metabolically coupled. Metabolic dysfunction of the RPE/Cho contributes to photoreceptor death in age-related macular degeneration (AMD). This study aimed to determine how aging, a major risk factor for AMD, alters the proteome and metabolic pathways in the retina and RPE/Cho.

**Methods:** We performed quantitative proteomics on neural retina and RPE/Cho from young (8-week) and old (73-week) C57BL/6J mice. The differentially expressed proteins between old and young, as well as between the retina and RPE/Cho, were visualized using volcano plots and heatmaps, and functional analyses were assessed by Gene Ontology enrichment tools.

**Results:** Aging altered the expression of 85 retinal proteins and 272 RPE/Cho, with 16 shared between tissues. In the aged retina, proteins associated with NADP(H) metabolism, neurodegeneration, and immunity were mostly upregulated, while proteins in lipid metabolism, transcription, and epigenetic regulation decreased. In the aged RPE/Cho, all 21 significantly altered lysosomal enzymes and transport proteins increased, together with complement components, and photoreceptor outer segment proteins. These changes accompanied increases in proteins associated with lipid metabolism and protection against lipid peroxidation, while several glutathione metabolism enzymes decreased. Seven shared proteins involved in lipid and NADP(H) metabolism, changed in opposite directions between retina and RPE/Cho. Comparison of the retinal and RPE/Cho proteomes identified 3073 differentially abundant proteins, of which 2452 maintained tissue-specific differences across both age groups, while approximately 17% showed age-dependent shifts in tissue enrichment. These shifts involved proteins associated with metabolism, extracellular matrix, visual function, lysosomal pathways, and innate immunity. Within the metabolic proteome, several glycolytic and fatty acid oxidation proteins showed greater relative enrichment in RPE/Cho with age, whereas glucose transporters and mitochondrial complexes shifted toward the retina.

**Conclusions:** Overall, these findings show that aging differentially remodels the retina and RPE/Cho proteome, accompanied by altered metabolic specialization between the two tissues.

## Introduction

The neural retina is among the most metabolically demanding tissues in the body, with photoreceptors in the outer retina requiring extraordinary amounts of energy and biosynthetic substrates to sustain phototransduction, synaptic activity, and continuous outer segment renewal [1–3]. The avascular outer retina depends largely on the choroidal circulation for oxygen, glucose, and other essential nutrients, which must traverse the retinal pigment epithelium (RPE) before reaching photoreceptors [4–7]. In addition to regulating nutrient transport, the RPE supports photoreceptor function by recycling visual chromophores, phagocytosing shed outer segments, and clearing metabolic waste from photoreceptors. Healthy RPE can utilize various alternative fuels derived from photoreceptors or choroid circulation, including lactate, succinate, proline, glutamine, and ketone bodies, to spare glucose for the photoreceptors [1,7–9]. Both RPE metabolic dysfunction and reduced choroidal blood flow have been implicated in retinal degeneration, particularly in age-related macular degeneration (AMD) [10–12].

Aging, the primary risk factor for AMD, causes progressive structural and functional decline in the retina, RPE, and choroid [13–17]. Photoreceptor density and retinal thickness, particularly in the macular region, decrease with age [13,14,18,19], along with impaired dark adaptation and reduced visual contrast sensitivity [20–22]. In both humans and mice, the RPE undergoes morphological disorganization [17,23], progressive loss of melanin [24,25], accumulation of lipofuscin [26,27], and reduced elasticity of the Bruch’s membrane-choroid complex [28] with age. With age, the choroidal vasculature thins and choriocapillaris flow density decreases, which may impair nutrient transport to the RPE and outer retina [29–31].

Early metabolic dysfunction can be an important contributor to age-related structural and functional decline in the retina and RPE/Cho. Transcriptomics of mouse retina identified reduced expression of genes involved in glycolysis and ATP synthesis in rod photoreceptors during early (3-12 months, equivalent to 20-40 years in humans) and intermediate (12-18 months, equivalent to 40-61 years in humans) ages [32]. Compared with young primates, aged primates (approximately 14 years old) showed a greater than 70% reduction in retinal ATP content and decreased cytochrome c oxidase activity [33]. Our previous metabolomic study found that glycolytic intermediates decreased in the retina but accumulated in RPE/Cho of intermediate-age mice (73-week) [34]. *Ex vivo* isotope tracing further demonstrated reduced glutamine use for TCA cycle anaplerosis in aged RPE/Cho, despite largely preserved glucose metabolism [35]. Genome-wide profiling of aging mouse rods also identified DNA methylation changes associated with energy metabolism [36]. Despite these transcriptomic and metabolomic findings, proteomic studies of retinal aging have largely examined the retina, RPE, or choroid separately, often across different ages and species [37–41]. Because the retina and RPE are metabolically interdependent, how aging remodels their proteomes in both tissues, particularly proteins involved in metabolism, remains poorly understood. Furthermore, loss of proteostasis and altered intercellular communication are established hallmarks of aging [42,43]. In this study, we used quantitative proteomics to identify age-related proteomic signature and evaluate changes in metabolic coupling between the neural retina and RPE/Cho in young and old (intermediate-aged) mice.

## Material and Methods

### Reagents

All reagents and resources used in this study are detailed in the Key Resources Table (**Table S1**).

### Animals

Seven-week-old (young) and 72-week-old (old) wild-type (WT) female C57BL/6J mice were purchased from the Jackson Laboratory and acclimated at West Virginia University for one week prior to experiments. All mouse experiments were performed in accordance with the National Institutes of Health guidelines and were approved by the Institutional Animal Care and Use Committee of West Virginia University.

### Sample collection and preparation

Mice were carried to the procedure room and acclimated for at least 1 hour before tissue collection. To minimize the effects of photoreceptor outer segment phagocytosis by the RPE, mice were euthanized between 2:00 and 4:00 PM under ambient light [44,45]. Following cervical dislocation, extraocular soft tissues, including muscle and fat, and the anterior parts were removed. Neural retina and RPE/Cho tissues were then rapidly isolated. Tissues were lysed in RIPA buffer supplemented with protease and phosphatase inhibitors (5mg/mL), and protein extracts were clarified by centrifugation at 12,000 rpm at 4 °C. Supernatants containing proteins were collected and samples were shipped to IDeA National Resource (Little Rock, AR) for quantitative proteomic analysis as reported [46].

### Quantitative proteomics

Proteins were reduced, alkylated, and purified by chloroform/methanol extraction prior to digestion with sequencing grade modified porcine trypsin (Promega). Tryptic peptides were then separated by reverse phase XSelect CSH C18 2.5 µm resin (Waters) on an in-line 150 x 0.075 mm column using an UltiMate 3000 RSLCnano system (Thermo). Peptides were eluted using a 60 min gradient from 98:2 to 65:35 buffer A:B ratio (Buffer A: 0.1% formic acid, 0.5% acetonitrile; Buffer B: 0.1% formic acid, 99.9% acetonitrile). Eluted peptides were ionized by electrospray (2.2 kV) followed by mass spectrometric analysis on an Orbitrap Exploris 480 mass spectrometer (Thermo). To assemble a chromatogram library, six gas-phase fractions were acquired on the Orbitrap Exploris with 4 m/z data-independent acquisition (DIA) spectra (4 m/z precursor isolation windows at 30,000 resolution, normalized AGC target 100%, maximum inject time 66 ms) using a staggered window pattern from narrow mass ranges using optimized window placements. Precursor spectra were acquired after each DIA duty cycle, spanning the m/z range of the gas-phase fraction (i.e. 496-602 m/z, 60,000 resolution, normalized AGC target 100%, maximum injection time 50 ms). For wide-window acquisitions, the Orbitrap Exploris was configured to acquire a precursor scan (385-1015 m/z, 60,000 resolution, normalized AGC target 100%, maximum injection time 50 ms) followed by 50x 12 m/z DIA spectra (12 m/z precursor isolation windows at 15,000 resolution, normalized AGC target 100%, maximum injection time 33 ms) using a staggered window pattern with optimized window placements. Precursor spectra were acquired after each DIA duty cycle.

### Proteomics data analysis

Following data acquisition, data were searched using an empirically corrected library and a quantitative analysis was performed to obtain a comprehensive proteomic profile. Proteins were identified and quantified using EncyclopeDIA [47] and visualized with Scaffold DIA using 1% false discovery thresholds at both the protein and peptide level. Protein exclusive intensity values were assessed for quality using an in-house ProteiNorm app, a tool for systematic evaluation of normalization methods, imputation of missing values and comparisons of multiple differential abundance methods [48]. Normalization methods evaluated included log2 normalization (Log2), median normalization (Median), mean normalization (Mean), variance stabilizing normalization (VSN) [49], quantile normalization (Quantile) [50], cyclic loess normalization (Cyclic Loess) [51], global robust linear regression normalization (RLR) [52], and global intensity normalization (Global Intensity) [52]. The individual performance of each method was evaluated by comparing of the following metrices: total intensity, pooled intragroup coefficient of variation (PCV), pooled intragroup median absolute deviation (PMAD), pooled intragroup estimate of variance (PEV), intragroup correlation, sample correlation heatmap (Pearson), and log2-ratio distributions.

### Proteomic pathway analysis

Bioinformatics analyses were performed on EnrichR and DAVID Bioinformatics [53]. Functional enrichment of differentially expressed proteins was assessed using Gene Ontology biological process (GOBP), cellular component (GOCC), and molecular function (GOMF), as well as Kyoto Encyclopedia of Genes and Genomes (KEGG) pathways and UniProt annotations. Metabolic proteins were identified by matching the proteomic dataset to our previously reported metabolic gene list [54], followed by functional annotation of the identified proteins.

### Statistics

Normalized proteomic data were analyzed using linear models for microarray data (limma), with empirical Bayes (eBayes) moderation of the standard errors [51]. To assess age-related shifts in protein abundance between retina and RPE/Cho, the difference between the old and young retina-to-RPE/Cho log-FC values was calculated (ΔDF = DF_old − DF_young). A cutoff of |ΔDF| ≥ 0.8 was used to define age-affected proteins. This threshold corresponds to a ∼1.74-fold change in the relative retina-to-RPE/Cho abundance ratio, which balances sensitivity with the ratio compression.

## Results

### Aging alters the retinal proteome

To investigate the impact of aging on the retinal proteome, we performed quantitative proteomic analysis of neural retina and RPE/Cho from young and old mice using DIA mass spectrometry. Given the functional and pathological interdependence of these tissues [1,55–57], age-related proteomic changes were analyzed separately in retina and RPE/Cho, followed by comparisons between the two tissues (**Figure 1**).

**Figure 1.**
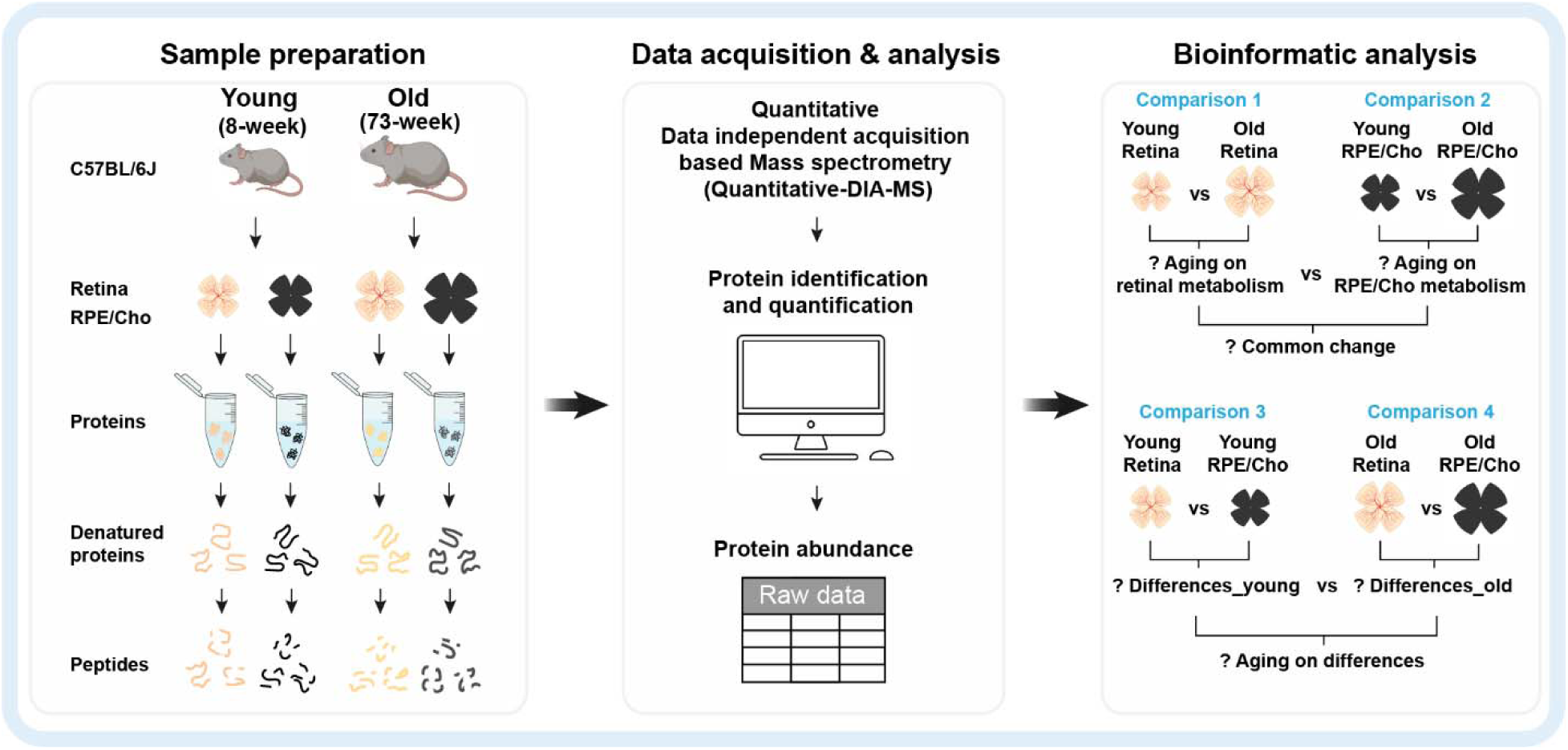
Quantitative proteomics workflow. Retina and RPE/Cho tissues were collected from young (8-week) and old (73-week) C57BL/6J mice and processed for quantitative DIA mass spectrometry-based proteomics. Following protein identification and quantification, proteomic profiles were analyzed using four comparisons: age-dependent changes in the retina (comparison 1), age-dependent changes in the RPE/Cho (comparison 2), tissue-specific differences between retina and RPE/Cho in young (comparison 3) and old (comparison 4), and common or age-specific changes across tissues. (N=5)

In the neural retinas, we detected a total of 4789 proteins with 46 proteins (0.94%) upregulated and 39 (0.80%) downregulated with age (**Figure 2A-B; Table S2**). Functional classification of the significantly altered proteins based on GOBP annotations and literature identified seven major categories, including metabolism, transcription, immunity, extracellular matrix (ECM) and focal adhesion, cytoskeleton, and neurodegeneration (**Figure 2C; Table S3**). Metabolic proteins represented the largest group (29%), followed by transcription (14%) and cytoskeleton (10%) (**Figure 2C**). Further GO analysis showed that proteins increased with age were enriched in immune-related processes, including IgE binding and major histocompatibility complex class I (MHC 1)-associated functions. In contrast, proteins decreased with age were associated with lipid metabolism, transcriptional regulation, and nuclear localization (**Figure S1; Table S4**). KEGG pathway analysis highlighted metabolism among the major pathways associated with altered proteins in the retina, followed by peroxisome proliferator-activated receptor (PPAR) signaling, drug metabolism, antigen processing, and selenocompound metabolism (**Figure 2D; Table S5)**.

**Figure 2.**
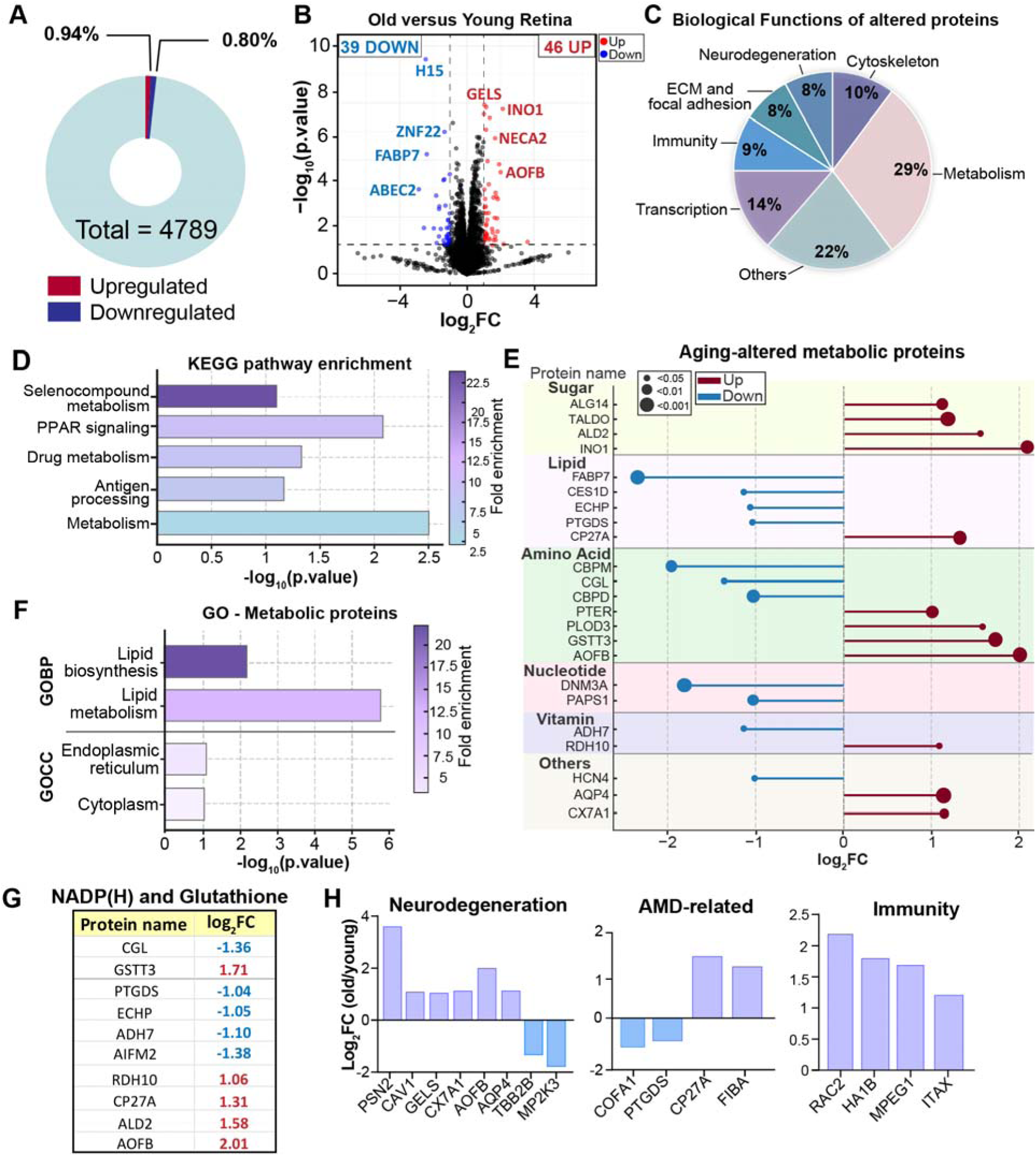
Aging alters the retinal proteomic profile. **(A)** The percentage of altered proteins identified in the retina between old and young mice (|log-FC| > 1, p < 0.05, N=5). **(B)** Volcano plot of changed protein in old versus young retina. **(C)** Classification of significantly altered proteins into 7 functional categories based on UniProt biological functions. **(D)** KEGG pathway enrichment analysis of significantly altered proteins. **(E)** Significantly altered metabolic proteins grouped by major metabolic pathways. **(F)** Gene Ontology (GO) enrichment analysis of altered metabolic proteins for biological processes (GOBP) and cellular components (GOCC). **(G)** Significantly altered NADP(H)- and glutathione-related metabolic proteins in old versus young retina. **(H)** Aging-associated changes in proteins linked to neurodegeneration, age-related macular degeneration (AMD), and immunity. Extracellular matrix (ECM); Peroxisome Proliferator-Activated Receptor (PPAR); reduced Nicotinamide Adenine Dinucleotide Phosphate (NADPH).

Based on metabolic substrates, we found proteins involved in amino acid, lipid, and sugar metabolism were mostly affected with age. GO analysis of the metabolic proteins identified lipid metabolism and lipid biosynthesis among the enriched biological processes, while cellular component analysis showed enrichment in the endoplasmic reticulum and cytoplasm (**Figure 2F; Table S6**). NADP(H)-related proteins, including ADH7, AIFM2, RDH10, CP27A, ALD2, AOFB, and glutathione-related proteins, including CGL and GSTT3, were significantly enriched with age.

Interestingly, eight proteins were related to neurodegeneration (Alzheimer’s and Parkinson’s disease), including PSN2, CAV1, GELS, CX7A1, AOFB, AQP4, TBB2B, and MP2K3, and six of them were upregulated in aged retina. We also found proteins implicated in AMD (COFA1, PTGDS, CP27A, FIBA) were altered in the aged retina, while all four differentially abundant immune proteins (RAC2, HA1B, MPEG1, ITAX) were upregulated (**Figure 2H**). In summary, these data demonstrate that aging alters the abundance of retinal proteins in key metabolic pathways, including lipid, amino acid, and sugar metabolism, with notable changes in NAD(P)H- and glutathione-related proteins, as well as proteins associated with neurodegeneration, AMD, and immunity.

### Aging alters the RPE/Cho proteome

Compared with the neural retina, RPE/Cho showed more extensive age-related changes, with 134 proteins upregulated (2.79%) and 138 downregulated (2.88%) (**Figure 3A, S2A; Table S7**). GO analysis of upregulated proteins identified enrichment of functions such as lipid metabolism, innate immunity, and vision-related processes (**Figure S2B; Table S8**). These proteins were associated with lysosomes, endoplasmic reticulum, endosomes, and cellular stress responses, with molecular functions including hydrolase, glucosidase, oxidoreductase, serine protease, and immunoglobulin E (IgE) binding activities. In contrast, downregulated proteins were associated with mRNA processing, cell adhesion, and stress-response pathways and were enriched in nuclear, ECM, and extracellular compartments. The molecular functions of the downregulated proteins included ribonucleoproteins, DNA-binding, ribosomal proteins, and chaperones (**Figure S2C-D; Tables S8 and S9**). Approximately 24% of the 272 significantly altered proteins were involved in metabolism, including lipid, sugar, amino acid and peptide, and nucleotide metabolism, as well as ion transport (**Figure 3B; Table S10**). Functional enrichment analysis further identified changes in lipid metabolism, including fatty acid, phospholipid, and prostaglandin metabolism. Cellular component analysis showed the strongest enrichment for lysosomal proteins, followed by proteins associated with the endoplasmic reticulum and peroxisomes (**Figure 3C; Table S11**). Notably, all altered lysosomal proteins were upregulated, including enzymes involved in sugar/glycan degradation (e.g., LYZ2, BGLR, GALNS, HPSE, MANBA, HEXA/B, and BGAL), lipid degradation (PLD3, ASAH1, and NAAA), proteolysis (CATZ, CATB, and CATS), and transport proteins (S15A4, CLCN7, NPC1, LAMP1, P2RX4, and CD68) (**Figure 3D**). Aging also affected proteins involved in redox regulation and lipid metabolism. GSTA3 and several NAD(P)H-related proteins (PTGR1, ADH7, XDH, and ALD2) increased, whereas GGT1, GPX7, and FMO2 decreased. Proteins associated with lipid metabolism and protection against lipid peroxidation (MBOA5, AIFM2, DHCR7, PLPL8, and FABP5) also increased, alongside phototransduction proteins (ARRS, RK, GNAT1, R9BP, and RHO) (Figure 3E). All significantly altered immune-associated proteins were upregulated, including complement factor H (CFAH), a genetic risk factor for AMD, and complement components C1QC, CO5, and C1QB (**Figure 3E**). Together, these findings suggest aging enhances oxidative stress, lipid peroxidation, and inflammation with impaired lysosomal processing, which might contribute to age-related functional decline (**Figure 3F**).

**Figure 3.**
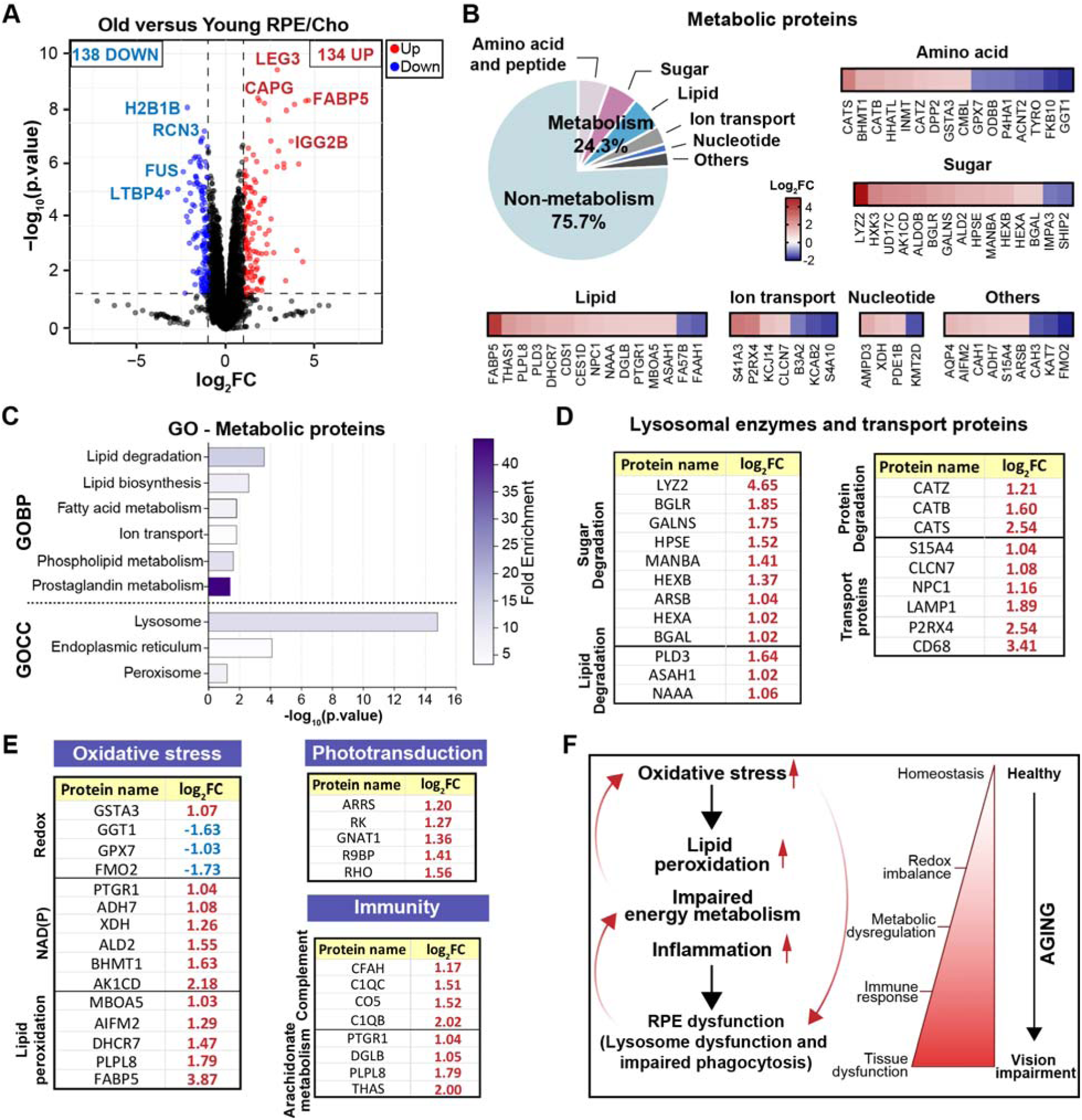
Aging alters the RPE/Cho proteomic profile. **(A)** Volcano plot of altered proteins in old versus young RPE/Cho. (|log-FC| > 1, p < 0.05, N=5). **(B)** Functional classification of significantly altered proteins. Heatmaps show log-FC values for metabolic proteins grouped by pathways. **(C-E)** GO analysis of significantly altered metabolic proteins for GOBP, GOCC, localization, and functions. **(F)** Proposed model of aging-associated proteomic changes in RPE dysfunction and vision impairment. Nicotinamide Adenine Dinucleotide Phosphate (NADP).

### Aging differentially alters protein abundance, shifting the proteomic balance between the retina and RPE/Cho

Aging altered 272 proteins in RPE/Cho and 85 proteins in the retina, with only 16 proteins shared between the two tissues (**Figure S3A; Table S12**). Among the shared proteins, seven involved in NADP(H)-dependent and lipid metabolism (AIFM2, CES1D, ADH7, PTGDS/PTGR1, and FABP7/FABP5) changed in opposite directions between retina and RPE/Cho (**Figure S3B**), highlighting the tissue-specific effects of aging. To evaluate how aging alters tissue-specific proteins in the retina and RPE/Cho, we compared protein abundance between the two tissues in young and old mice. Proteins with established functions in the visual cycle, melanin biosynthesis, and membrane transport showed the expected tissue enrichment, confirming the tissue identity of our proteomic datasets (**Figure 4A**). Comparison of the retina with RPE/Cho proteome yielded 2731 differentially expressed proteins in young mice and 2794 in old mice, with 2452 proteins shared across age groups (**Figure 4B-D; Tables S13-15**).

**Figure 4.**
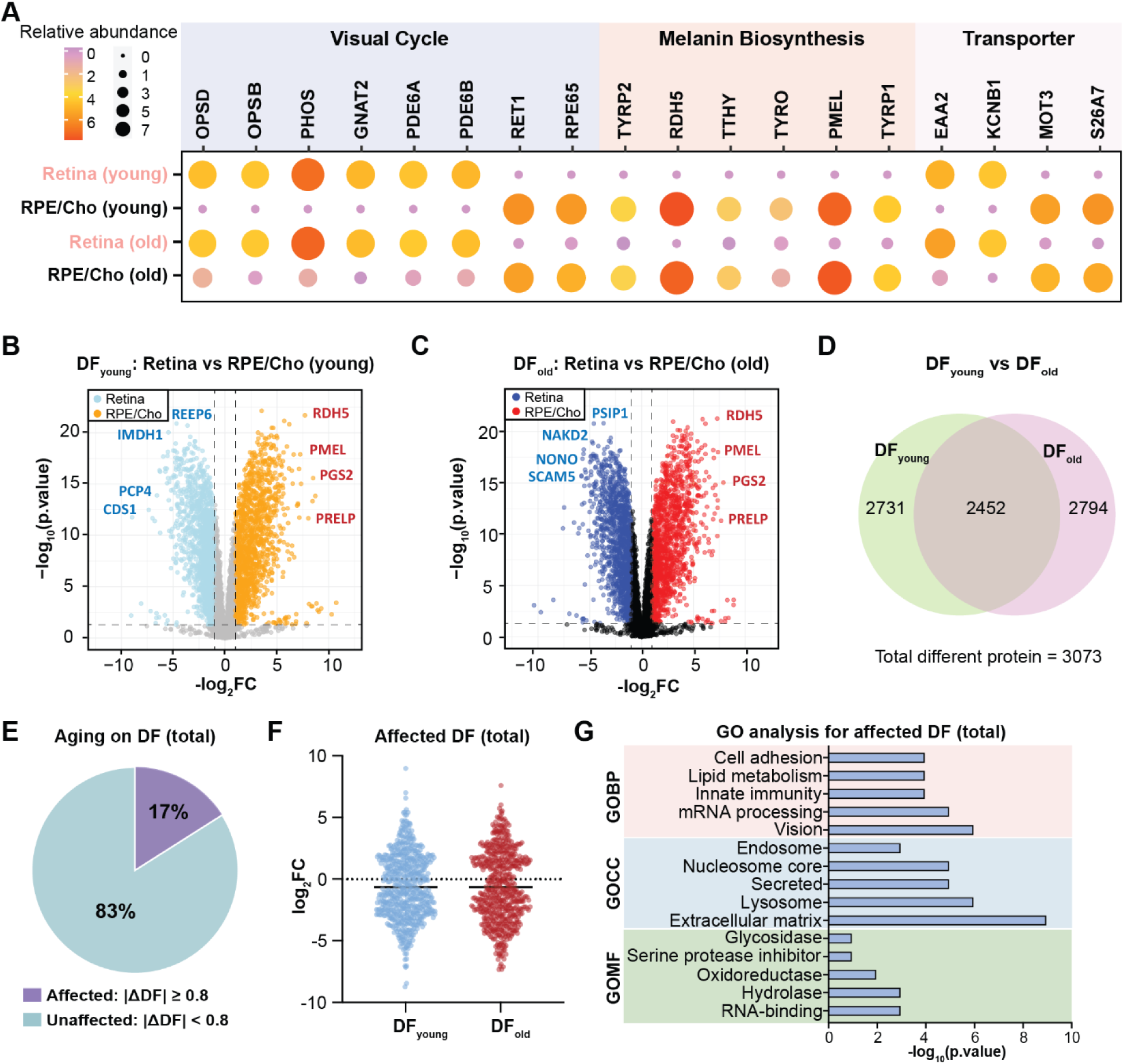
Aging alters proteomic differences between retina and RPE/Cho. **(A)** Relative abundance of protein markers in retina and RPE/Cho in young and old mice, normalized to young retina. **(B-C)** Volcano plots of protein difference in the retina versus RPE/Cho (DF) in young **(B)** and old **(C)** mice. Log-FC was calculated as retina vs RPE/Cho and plotted as log-FC, with negative values indicating higher expression in retina and positive values indicating higher expression in RPE/Cho (|log-FC| > 1, p < 0.05). **(D)** Venn diagram of DF proteins in young and old mice. **(E)** Proportion of age-affected DF proteins young and old mice (total = 3073 proteins). Proteins were defined as age-affected when the change in retina versus RPE/Cho log-FC between young and old mice (|ΔDF|) was ≥ 0.8. **(F)** Distribution of retina versus RPE/Cho log-FC values for differentially abundant proteins in young and old mice. **(G)** GO enrichment analysis of age-affected proteins for GOBP, GOCC, and GOMF terms. Mean ± SD; N=5.

To identify proteins with age-dependent shifts in tissue enrichment while minimizing false positives from multi-condition variance, we applied a log-fold change shift threshold of |ΔDF| ≥ 0.8 (∼1.74-fold) in the retina-to-RPE/Cho abundance ratio. Based on this criterion, approximately 17% of the proteins were affected by aging, whereas 83% maintained stable (**Figure 4E; Table S16**). Aging widened the distribution of log-FC values in both directions, higher in retina (positive) and/or higher in RPE/Cho (negative), demonstrating increased enrichment of proteins with aging between the two tissues (**Figure 4F; Table S16**). GO analysis showed that age-affected proteins were enriched in visual function, mRNA processing, innate immunity, lipid metabolism, and cell adhesion (**Figure 4G; Table S17**). These proteins were localized to the ECM, lysosome, secreted proteins, nucleosome core, and endosome, with molecular functions including RNA-binding, hydrolase, oxidoreductase, serine protease inhibitor, and glycosidase activities (**Figure 4G**). Together, these data demonstrate that most tissue differences were maintained during aging, while a subset became more pronounced, particularly among proteins associated with visual, immune, and metabolic functions.

### Aging differentially alters metabolic proteins between the retina and RPE/Cho

Among the 3073 differentially expressed proteins between retina and RPE/Cho, 769 were annotated as metabolic proteins (**Figure S4A; Table S18**). Specifically, 692 and 689 metabolic proteins differed between retina and RPE/Cho in young mice and old mice, respectively, with 612 shared between age groups (**Figure S4B; Tables S18-19**). These proteins were primarily associated with mitochondria, the nucleus, endoplasmic reticulum, and lysosomes (**Figure S4C**). Functional classification showed that pathways in amino acid and peptide, lipid, nutrient transport, sugar/glycan, and nucleotide metabolism accounted for most metabolic protein differences between retina and RPE/Cho (**Figure 5A; Table S18**). Proteins involved in the electron transport chain (ETC), glycolysis, the TCA cycle, nucleotide metabolism, and histone modification were predominantly enriched in the retina. In contrast, proteins involved in amino acid and peptide, lipid, sugar/glycan, redox, sulfate, and detoxification pathways were predominantly enriched in RPE/Cho. Transport and vitamin metabolism proteins showed a more mixed distribution (**Figure 5B**). These patterns are consistent with previously reported tissue-specific metabolic specialization of these two tissues [58,59].

**Figure 5.**
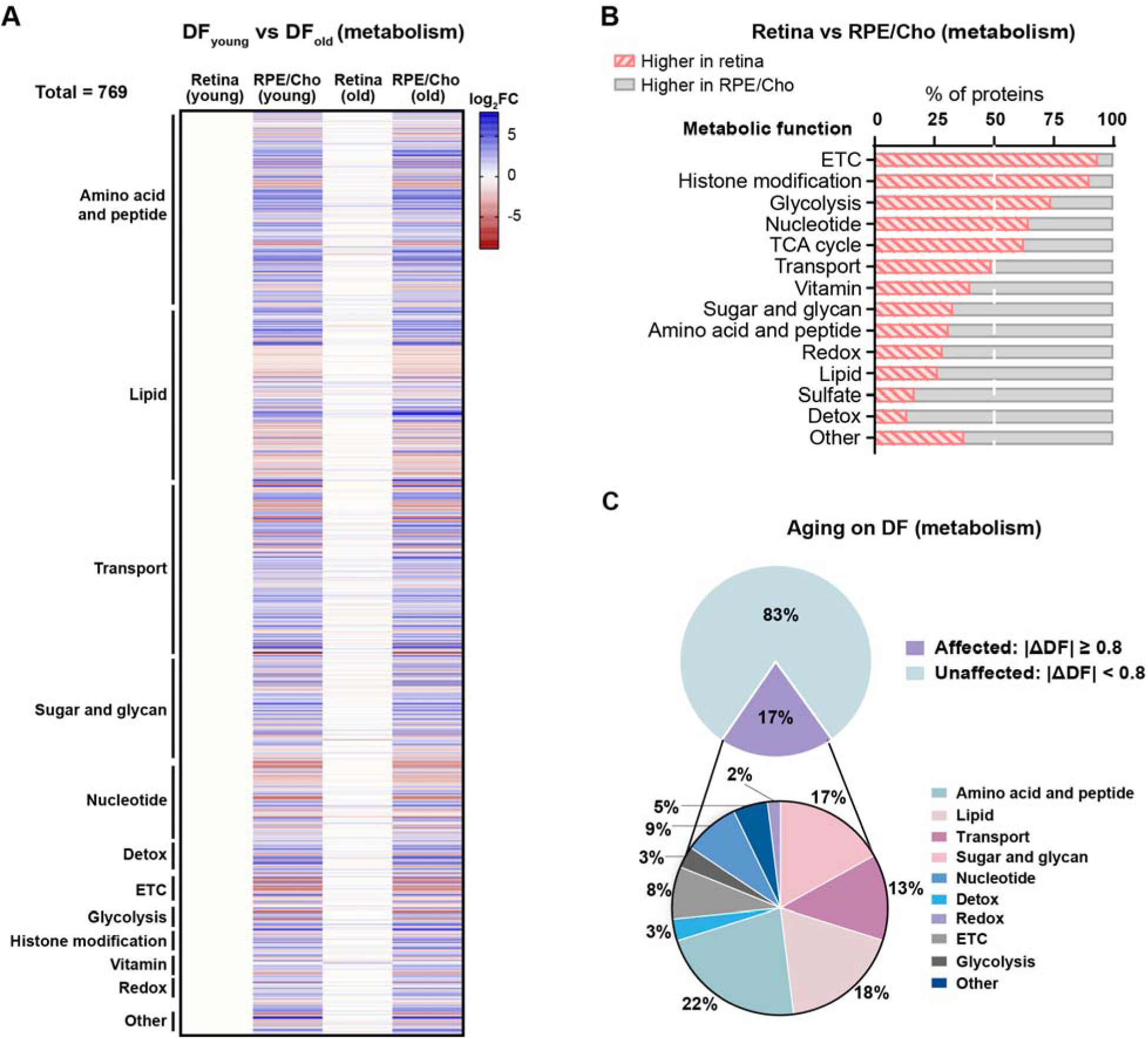
Aging alters metabolic differences between retina and RPE/Cho. **(A)** Heatmap of altered DF metabolic proteins in young and old mice, grouped by metabolic category. **(B)** Distribution of DF metabolic proteins in pathways. **(C)** Proportion and classification of DF metabolic proteins affected by aging (|ΔDF| ≥ 0.8). Electron transport chain (ETC).

Most tissue-specific differences in metabolic proteins were maintained with age, whereas approximately 17% of the proteins shifted their tissue distribution (**Figure 5C; Table S20**). Among the age-affected metabolic proteins, amino acid and peptide metabolism represented the largest group (22%), followed by lipid metabolism (18%), sugar and glycan metabolism (17%), and transport (13%). Smaller groups included proteins involved in nucleotide metabolism, the ETC, glycolysis, detoxification, redox, and vitamin metabolism (**Figure 5C**). Together, these data show that while retina and RPE/Cho preserve their overall metabolic specialization with age, aging selectively remodels a subset of metabolic and nutrient transport proteins.

### Aging alters metabolic specialization between retina and RPE/Cho

To determine how aging impacts metabolic specialization between retina and RPE/Cho, we compared age-affected metabolic proteins between the two tissues. These proteins were predominantly enriched in RPE/Cho (negative log_2_FC) at both ages, with a modest shift toward greater RPE/Cho abundance in old mice (**Figure 6A; Table S20**). Given the tight metabolic interdependence between the retina and RPE/Cho, we focused on energy metabolism pathways, including glucose metabolism, the ETC, fatty acid oxidation, pentose phosphate pathway (PPP), and NAD-metabolism (**Figure 6B; Table S20**). Key proteins in these pathways showed strong age-dependent shifts in their relative tissue distributions. Among ETC components, tissue differences in CIA30 and COX42 decreased with age, whereas NDUA3 and CX7A1 increased and shifted toward higher RPE/Cho enrichment. Proteins in glucose metabolism demonstrated variable shifts: tissue differences in glucose and monocarboxylate transporters (G6PT3, GTR1, and MOT4) narrowed with age, whereas the glycolytic enzymes (ALDOB and HXK3) became more enriched in RPE/Cho, while KPYM remained unchanged. Selected fatty acid metabolism proteins also trended towards greater RPE/Cho enrichment with age. CPT1B, a rate-limiting enzyme in mitochondrial fatty acid oxidation, was enriched in young RPE/Cho, and this difference increased with age. However, ACS2L, an enzyme involved in short-chain fatty acid activation, shifted from a small tissue difference to pronounced RPE/Cho enrichment. Conversely, PPP proteins showed minor alterations, and the tissue difference in the NAD-related protein NQO1 decreased with age (**Figure 6B**). Together, these findings suggest that aging rewires metabolic specialization between retina and RPE/Cho, particularly in glucose and fatty acid metabolism.

**Figure 6.**
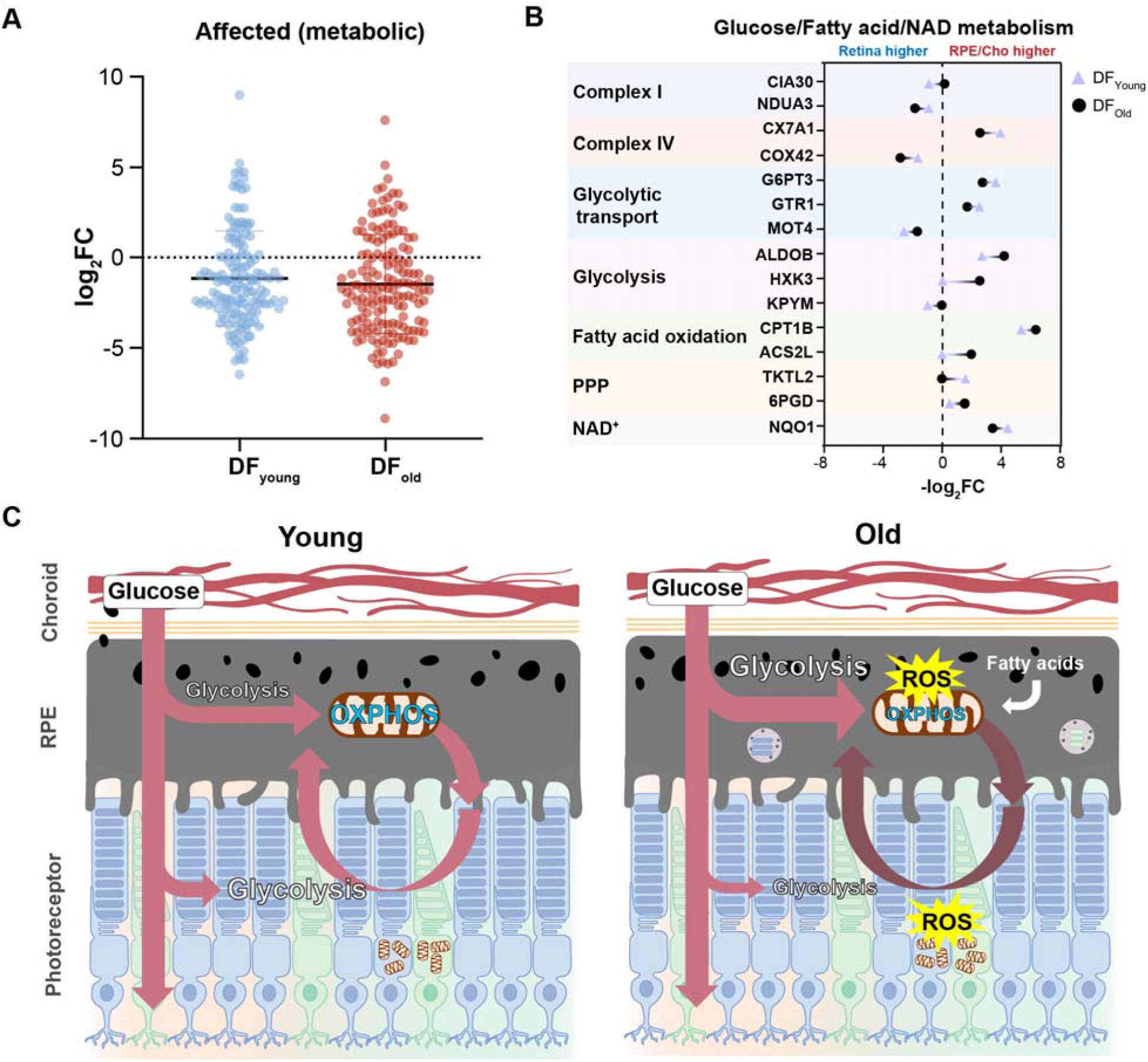
Aging remodels retina and RPE/Cho metabolism. **(A)** Distribution of age-affected metabolic DF proteins in young and old mice. Aging-affected DF proteins were defined by a change in the retina-to-RPE/Cho difference of |ΔDF| ≥ 0.8. **(B)** Shifts of metabolic pathways in aged retina and RPE/Cho. Triangles represent young mice and circles represent old mice; negative values indicate higher abundance in retina and positive values indicate higher abundance in RPE/Cho. **(C)** Schematic model of remodeled glucose metabolism in aged retina and RPE. Black granules represent RPE melanosomes, and brown structures represent mitochondria, where oxidative phosphorylation (OXPHOS) occurs. In aged RPE, small vesicles represent lysosomes containing accumulated or incompletely degraded material. The model depicts increased oxidative stress (ROS) affecting mitochondrial OXPHOS in both RPE and photoreceptors, with a possible shift toward greater RPE glycolysis that could reduce glucose availability to photoreceptors. Mean ± SD; N=5. Pentose phosphate pathway (PPP); Nicotinamide adenine dinucleotide (NAD^+^).

## Discussion

In this study, we found that aging alters the neural retina and RPE/Cho proteomes in a tissue-specific manner, with more extensive changes in the RPE/Cho. In the aged retina, these changes involved proteins associated with lipid and sugar metabolism, NADPH and glutathione metabolism, neurodegeneration, and immune responses. In contrast, the aged RPE/Cho showed increased abundance of lysosomal enzymes and transport proteins, photoreceptor outer segment proteins, and complement components, as well as changes in lipid metabolism and redox-related proteins. Most tissue-specific metabolic differences were maintained with age, whereas glycolytic and fatty acid utilization proteins shifted toward greater relative enrichment in RPE/Cho (**Figure 6B-C**). These findings indicate more extensive age-related remodeling in RPE/Cho, which may affect its metabolic support of the neural retina and contribute to retinal functional and structural decline.

The neural retina and RPE/Cho form an interdependent metabolic ecosystem. Glucose from the choroidal circulation passes through the RPE to reach photoreceptors, which convert much of it to lactate through aerobic glycolysis [1,4,7,60]. The RPE uses photoreceptor-derived lactate and other substrates to support mitochondrial metabolism, helping spare glucose for the outer retina. Besides glycolysis, glucose supports multiple essential pathways, including (1) oxidative phosphorylation for ATP production; (2) the PPP for NADPH and nucleotide synthesis; (3) myo-inositol synthesis; (4) the polyol pathway for sorbitol and fructose synthesis; (5) de novo lipid synthesis; (4) non-essential amino acid synthesis; and (7) protein and lipid glycosylation. Our data suggest that aging alters multiple pathways in glucose metabolism that may affect the metabolic ecosystem between retina and RPE/Cho.

Aging substantially altered the relative tissue enrichment of glycolytic enzymes, including hexokinase 3 (HXK3/HK3), aldolase B (ALDOB), and pyruvate kinase (KPYM/PKM). Tissue differences in glucose transporter 1 (GTR1/GLUT1), the glucose-6-phosphate transporter G6PT3, and monocarboxylate transporter 4 (MOT4/MCT4) also narrowed with age (**Figure 6B**). Consistent with this, our previous targeted metabolomics study of mice at the same ages (73-versus 6-week-old) found that glucose and glycolytic intermediates accumulated in aged RPE/Cho, whereas several glycolytic intermediates decreased in the retina [34]. Similarly, increased PKM2 expression and pyruvate kinase activity have been reported in aged mouse RPE [61], supporting age-related changes in glycolytic regulation. Increased RPE glycolysis has also been reported in models of AMD-like degeneration driven by defective mitochondrial respiration and mTOR activation [62] or by HIF stabilization [11,63]. In cultured human RPE, inhibition of mitochondrial respiration increases glucose consumption and glycolysis [64]. In line with these findings, we observed an age-related shift toward lower relative abundance of respiratory chain proteins in the RPE/Cho compared to the retina. Reducing GLUT1 expression in the RPE also lowers retinal glucose availability and induces stress in photoreceptors and Müller glia [65]. Together, these observations suggest that age-related changes in RPE/Cho glucose utilization and transport may limit glucose delivery to the outer retina and contribute to photoreceptor dysfunction and degeneration in aging and AMD.

Myo-inositol-1-phosphate synthase (INO1), which catalyzes the first step in myo-inositol synthesis from glucose-6-phosphate, was among the most strongly increased proteins in the aged retina. Myo-inositol is an abundant ocular osmolyte and a precursor of phosphoinositides, which regulate protein trafficking and phototransduction in the retina [66–68]. Elevated brain myo-inositol has been associated with aging and Alzheimer’s disease and is commonly interpreted as a marker of glial proliferation [69–71]. In our dataset, aging also altered Müller glia-associated proteins, including aquaporin-4 (AQP4) and fatty acid-binding protein 7 (FABP7) [72,73], together with proteins implicated in neurodegeneration, including presenilin-2 (PSEN2) [74,75], gelsolin (GSN) [76], and caveolin-1 (CAV1) [77,78]. These observations suggest that altered myo-inositol metabolism is linked to Müller glial remodeling and neurodegenerative stress in the aging retina.

A striking feature of our data is the alteration of NADP(H)-dependent redox metabolism. NADPH is the principal source of reducing power for antioxidant defense, lipid biosynthesis, and the reduction of all-trans-retinal for visual pigment recycling. Both glucose and glutamine can support NADPH generation in rod photoreceptors [79]. NADPH also maintains glutathione and thioredoxin in their reduced forms, supporting cellular antioxidant defenses. Aging affected multiple proteins involved in these systems. In retina, CGL, which supplies cysteine for GSH synthesis, decreased, whereas GSTT3, which detoxifies reactive electrophiles, and the hydrogen peroxide-producing enzyme MAOB increased. In RPE/Cho, GGT1, which supports GSH salvage, and the antioxidant enzymes GPX7 and FMO2 decreased, while GSTA3 and the ROS-generating enzyme XDH increased. Together, these changes indicate tissue-specific remodeling of redox-related pathways that may affect antioxidant capacity and response to oxidative stress in the aging retina and RPE/Cho.

NADPH is also required for fatty acid and cholesterol biosynthesis. In aged RPE/Cho, increased DHCR7 abundance was accompanied by increases in MBOA5, FABP5, PLPL8, and AIFM2, indicating changes in proteins involved in cholesterol synthesis, phospholipid remodeling, and lipid metabolism. DHCR7 requires NADPH to convert 7-dehydrocholesterol to cholesterol, whereas MBOA5/LPCAT3 incorporates polyunsaturated fatty acids into membrane phospholipids and can influence their susceptibility to peroxidation [80,81]. The cholesterol precursor 7-dehydrocholesterol is highly prone to peroxidation, and its oxidation products drive retinal degeneration in Smith-Lemli-Opitz syndrome [82,83]. The aged RPE may therefore face competing demands for NADPH to support lipid synthesis and protect against lipid peroxidation.

The aldo-keto reductase ALD2 was among the few proteins increased in both the aged retina and RPE/Cho. ALD2 belongs to the aldose reductase family, which can reduce glucose to sorbitol and fructose using NADPH in the polyol pathway [84]. Increased polyol flux can reduce NADPH availability and promote oxidative and osmotic stress, mechanisms implicated in diabetic retinopathy [85]. ALD2 can also reduce lipid-derived aldehydes, including 4-hydroxynonenal and cholesterol-derived aldehydes [84,86]. Thus, the increase of ALD2 and the related reductase AK1CD in RPE/Cho may reflect enhanced detoxification of lipid peroxidation products in addition to altered polyol metabolism [87]. Both processes increase NADPH demand, potentially limiting reducing equivalents available for glutathione recycling and lipid synthesis. Aldolase B (ALDOB) was also increased in RPE/Cho. Because ALDOB cleaves both fructose-1-phosphate and fructose-1,6-bisphosphate, its increase may indicate altered fructose utilization [88]. Future studies measuring sorbitol and fructose flux will help determine the relative contributions of these pathways in the aged retina and RPE/Cho.

Lysosomal remodeling was another prominent feature of aging in RPE/Cho. Nearly every major class of lysosomal hydrolases was increased, and the lysosome was the most strongly enriched cellular component among the upregulated proteins with age. The RPE is highly phagocytic, engulfing and degrading approximately 10% of photoreceptor outer segments daily [89,90]. In addition to hydrolases, several lysosomal membrane and transport proteins increased, including LAMP1, the cholesterol exporter NPC1, the Cl□/H□ antiporter CLCN7, the peptide/histidine transporter SLC15A4, and the lysosomal ATP-gated channel P2RX4. Many of these proteins are components of the transcription factor EB (TFEB)-regulated lysosomal gene network [91,92], suggesting expansion of lysosomal response to increasing degradative burden. However, increased lysosomal protein abundance does not necessarily indicate more efficient degradation. Elevated lysosomal pH can reduce hydrolase activity and impair outer-segment degradation, while accumulation of lipofuscin bisretinoids such as A2E can disrupt lysosomal processing [93,94]. Slower phagosome processing and reduced phagolysosome formation have been reported in aged mouse RPE [95]. In our dataset, increased abundance of photoreceptor proteins, including rhodopsin (RHO), GNAT1, and arrestin (ARRS), may reflect persistence of incompletely degraded outer-segment material. Additionally, aged subretinal microglia can engulf outer segments and accumulate lipofuscin-like material [96]. The marked increases in LYZ2 (∼25-fold) and CD68 (∼10.6-fold), proteins associated with macrophages and microglia [97–99], may therefore indicate a contribution from these immune cells to the lysosomal signature in aged RPE/Cho.

Overall, our findings support a model in which aging progressively uncouples the metabolic interdependence between the retina and RPE/Cho. Aged RPE/Cho shows increased abundance of glycolytic and lysosomal proteins, together with changes in proteins involved in mitochondrial respiration, glucose transport, and redox regulation. In contrast, the retina shows distinct changes in glucose-derived biosynthetic and antioxidant pathways. This remodeling may affect the ability of RPE/Cho to support the energetic and redox demands of the outer retina and could contribute to susceptibility to retinal degeneration and AMD.

## Acknowledgments

This work was supported by National Eye Institute Grant EY032462 (JD), EY039633 (JD), and the Retina Research Foundation (JD). The proteomics study was supported by IDeA National Resource for Quantitative Proteomics and NIH/NIGMS grant R24GM137786.

## Supporting Information

This article contains supporting information (**Supplementary Figures** and **Tables**).

## Author Contributions

Conceptualization, J.D; Investigation, S.Z., A.P., R.M., D.A., T.N., J.D.; Writing, S.Z., A.P., J.D.; Funding Acquisition J.D; Supervision, J.D.

## Conflicts of interest

The authors declare no conflicts of interest.

## Ethical approval

The mouse experiments were performed in accordance with the National Institutes of Health guidelines and the protocol approved by the Institutional Animal Care and Use Committee of West Virginia University.

## Informed consent

The authors declare no patients were involved in this study; therefore, no informed consent was required.

## Data Availability Statement

Proteomics data generated in this study were deposited in MassIVE (ID: MSV000097974).

## Abbreviations

RPE/Cho: Retinal pigmented epithelium/Choroid
AMD: Age-related macular degeneration
DIA: Data-independent acquisition
MS: Mass spectrometry
GO: Gene ontology
PPAR: Peroxisome proliferator-activated receptors
ETC: Electron transport chain
ECM: Extracellular matrix.

## Notes

### Competing Interest Statement

The authors have declared no competing interest.

